# Nanodisc reconstitution and characterization of amyloid-β precursor protein C99

**DOI:** 10.1101/2024.04.21.590446

**Authors:** Bankala Krishnarjuna, Gaurav Sharma, Volodymyr M Hiiuk, Jochem Struppe, Pavel Nagorny, Magdalena I Ivanova, Ayyalusamy Ramamoorthy

## Abstract

Amyloid precursor protein (APP) plays a pivotal role in the pathology of Alzheimer’s disease. Since the fragmentation of the membrane-bound APP that results in the production of amyloid-beta peptides is the starting point for amyloid toxicity in AD, it is important to investigate the structure and dynamics of APP in a near-native lipid-bilayer environment. However, the reconstitution of APP into a stable/suitable membrane-mimicking lipid environment is a challenging task. In this study, the 99-residue C-terminal domain of APP is successfully reconstituted into polymer nanodiscs and characterized using size-exclusion chromatography, mass spectrometry, solution NMR, and magic-angle spinning solid-state NMR. In addition, the feasibility of using lipid-solubilizing polymers for isolating and characterizing APP in native *E. coli* membrane environment is demonstrated.

## INTRODUCTION

Membrane proteins embedded in the cell membrane or anchored to the cell membrane play major roles in various cellular processes such as cell signaling, molecular transport, adhesion processes, and drug metabolism and, therefore, are primary targets for pharmaceutical drug development (1-3). Hence, it is essential to explore high-resolution structures of membrane proteins to understand the underlying molecular mechanism of their function and thus to assist drug design and development projects (4-6). However, the *in vitro* characterization of membrane proteins is challenging, as their isolation/purification and, importantly, their reconstitution into a stable/suitable membrane-mimicking lipid environment is difficult in general (7-10).

Although detergents are cost-effective and efficient in extracting membrane proteins, they have limitations, as their short alkyl chains and headgroups are chemically distinct from those of native membrane lipids, rendering them discrete physico-chemical properties that can affect the function of membrane proteins (5, 11-13). Moreover, their denaturing properties make it difficult to maintain membrane protein’s oligomeric or multimeric states that are stabilized by intermolecular interactions (14-17). Planar lipid-bilayer-containing bicelles have been used extensively to study the structure, dynamics, and membrane topology of a variety of membrane proteins (18). However, bicelles also contain detergent-like short-chain lipids that could affect the stability of a reconstituted membrane protein (19-21). Likewise, liposomes are also limited by a lack of good stability and vesicular morphology (9, 22, 23). The membrane-mimicking systems containing detergents provide minimal stability (13, 24), and may not be desirable for high-resolution membrane protein studies (25, 26). On the other hand, nanometer-sized lipid bilayered systems encircled and stabilized by a protective belt of peptides/proteins/polymers (nanodiscs) are shown to provide a stable and more native-like membrane environment for structural and functional characterization of a variety of membrane proteins (5, 27-37).

This study aims to demonstrate the feasibility of using nanodiscs to study the transmembrane protein domain C99 of the full-length amyloid precursor protein (APP). C99 plays important roles in the pathology of Alzheimer’s disease (AD). APP is an integral membrane protein with 695-770 amino acid residues (38). It is expressed in many different tissues, especially in the synapses of neurons, and its main biological function is yet to be understood (39, 40). However, APP is extensively studied in connection with its amyloid-β (Aβ) peptide, the key component of plaques in the pathogenesis of AD (41-44). APP is non-systematically cleaved by secretases; thus, Aβ peptide fragments with different lengths are produced within the cell membrane (45-47). The initial processing of APP by β-secretase results in a membrane-anchored C-terminal (99 residues) fragment (β-CTF) called C99 and a secreted fragment sAPPβ (38, 39). Further sequential processing of C99 by γ-secretase produces Aβ40 and Aβ42 peptide fragments within the cell membrane (38, 46). Previous studies have reported the importance of the lipid membrane and membrane composition on the structure and stability of APP and C99. Therefore, to gain a molecular-level understanding of the processes underlying the production of Aβ peptides and the roles of dynamic-structure and membrane interaction of APP, it is essential to study APP (or C99) in a near-native lipid membrane.

C99 was overexpressed in *E. coli* (BL21-C41strain) and solubilized using a nanodisc-forming non-ionic pentyl-inulin polymer (5, 48-50). The polymer-solubilized APP in *E. coli* native lipids was characterized by SDS-PAGE and Western blot. Protein was also purified under denaturing conditions and reconstituted into polymer nanodiscs (pentyl-inulin-based nanodiscs). The APP reconstituted in nanodiscs was characterized by ^1^H NMR spectroscopy, dynamic-light scattering (DLS), differential scanning calorimetry (DSC), circular dichroism (CD) spectroscopy, mass spectrometry, and transmission electron microscopy (TEM) experiments. A ^15^N-labelled C99 expressed and purified from *E. coli* was also reconstituted into inulin-based polymer nanodiscs and characterized by high-resolution magic angle spinning (MAS) solid-state NMR spectroscopy.

## EXPERIMENTAL SECTION

### Protein expression

The plasmid (T7 promotor) expressing APP with a 6xHis tag at the C-terminus (APP-C99) or APP with a 10xHis tag at the N-terminus (APP-NT2), was transformed into C41-competent cells (Lucigen Corporation, WI, USA). After 16 h at 37 °C, a single colony from the transformation plate was inoculated into 10 mL and grown overnight at 37 °C (starter culture). The protein was over-expressed by the autoinduction method as previously reported (51, 52). Briefly, the starter culture was inoculated into ZYM-5052 autoinduction media (ZY - 1% tryptone, 0.5% yeast extract; M - 0.05M KH_2_PO_4,_ 0.05M Na_2_HPO_4,_ and 0.025M (NH_4_)_2_SO_4,_ and 5052 –0.5% glycerol, 0.05% glucose, and 0.2% lactose (pH 6.75)) containing 1 mM MgSO_4_, 1x trace metal mix (Hollister, CA, USA), 0.1x Basal Medium Eagle (BME) vitamin (Sigma-Aldrich), and 100 µg/mL ampicillin and were grown at 37 °C until OD_600_ reached ∼0.8. Then the temperature was reduced to 20 °C and the overexpression was allowed for 20 h. The cells were pelleted down by centrifugation at 6500 xg for 10 min at 4 °C and stored at -80 °C until protein solubilization/purification. For preparing ^15^N-labeled C99 protein, ^15^NH_4_Cl was added to the buffers in the autoinduction media (1.5 g/L ^15^NH_4_Cl, 0.005M Na_2_SO_4,_ 0.05M KH_2_PO_4_ and 0.05M Na_2_HPO_4_).

### Cell lysis and membrane preparation

Cells were resuspended in 10 mM Tris buffer (pH 7.4) containing 100 mM NaCl, lysozyme (5 mg/mL) (New Brunswick, NJ, USA), DNase (1 mg/mL) (Sigma-Aldrich, St. Louis, Missouri, USA) and 2 mM MgCl_2_ (Sigma-Aldrich), and lysed by sonication using a 13 mm sonicator probe (Thomas Scientific, LLC, NJ) with 8 cycles of 10 s pulse and 1 min cooling between every 2 pulses. The soluble components were removed by centrifugation, and the membranes were collected and washed twice using the washing buffer (75 mM Tris pH 7.8, 300 mM NaCl, 0.2 mM EDTA). To remove the EDTA, a third wash was given with the resuspension buffer (10 mM Tris buffer, pH 7.4, and 100 mM NaCl). cOmplete™ protease inhibitor cocktail (Sigma-Aldrich) was included in all the steps of protein purification.

### Pentyl-inulin synthesis

Natural inulin polymer extracted from chicory roots was purchased from Sigma-Aldrich. Pentyl-inulin (average molecular weight of ∼3 kDa) polymer used in this study was chemically synthesized by functionalizing inulin with hydrophobic pentyl groups using pentyl bromide (Sigma-Aldrich), purified and characterized as described elsewhere (48).

### Membrane solubilization using pentyl-inulin

Cell membranes were weighed using an analytical balance and resuspended (25 mg/mL) in 10 mM Tris buffer (pH 7.4) containing 100 mM NaCl and mixed with polymer at a 1:1 w/w membrane-to-polymer ratio (53). The solubilization was carried out at 4, 25 and 37 °C. Then, the insoluble components were removed by centrifugation at 10,000 rpm, 4 °C for 45 min. The clear supernatant was collected carefully for further analysis. In addition, as a control, the membranes were also solubilized in the presence of 0.5 % n-dodecyl-beta-maltoside (DDM) detergent.

### Western blotting

SDS-PAGE was performed on the solubilized samples using 4-12% (gradient) bis-tris NuPAGE gel (Thermo Fisher Scientific, MA, USA). 1X MES NuPAGE was used as a gel running buffer. The gel was run at 100 V for 10 min and then 120 V for 70 min at RT. While running SDS-PAGE, the PVDF membrane (Immuno-Blot Bio-Rad 1620177, 26cm x 3.3m, 0.2um) was soaked in 100% methanol for 30-60 min, followed by soaking in the transfer buffer (0.25M Tris, 1.92M glycine and 20% methanol) for 5-10 min. Transfer from SDS-PAGE gel onto PVDF membrane was carried at 4 °C, 110 V for 1 h. Then, the PVDF membrane was blocked with the blocking buffer (10% milk in TBST buffer – 0.02M Tris (pH 7.6), 0.15M NaCl, and 0.1% Tween 20) for 1h at room temperature. The blot was probed with the primary antibody (6E10; Cat# 803004) (Bio Legend, San Diego, California, USA; prepared at a dilution of 1:500 in 5% milk TBST) and incubated overnight at 4 °C with gentle shaking. The next day, the membrane was washed three times with 5% milk TBST before incubating with the secondary antibody (goat anti-mouse HRP, Thermo Fisher Scientific; Cat#32230; prepared at a dilution of 1:5000 in 5% milk TBST). The membrane was washed as previously, and the proteins were detected in the presence of substrate, Eco bright Femto HRP.

### Purification of APP-C99

The membranes were solubilized overnight in the solubilization buffer (20 mM Tris pH 7.8, 150 mM NaCl, 8M urea, 0.2% SDS) at 4 °C with gentle rotation. The insoluble material was removed by centrifugation at 39, 000 xg, 4 °C for 30 min. The supernatant was collected and filtered using a 1.2-micron cellulose acetate filter (Findlay, OH, USA). Then, the APP-C99 was purified by Ni-NTA affinity chromatography (Chicago, IL, USA). The sample was loaded onto a 5 mL Ni-NTA column equilibrated with the same solubilization buffer at a 2 mL/min flow rate using FPLC. Then, the column was washed using 50 mL of 10 mM Tris buffer (pH 7.4) containing 100 mM NaCl and 3 % Empigen BB (Sigma-Aldrich) to replace urea and SDS with Empigen. The APP-C99 was eluted from the column using an imidazole gradient (0-300 mM). The fractions from the elution peak (280 nm) were analyzed by SDS-PAGE (Piscataway, NJ, USA), and the fractions containing the APP-C99 protein were pooled. The concentration of APP-C99 was determined by UV spectroscopy using an extinction coefficient of 5960 M^-1^ cm^-1^.

### Preparation of liposomes and polymer nanodiscs

7 mg of DMPC and 3 mg of DMPG (Avanti Polar Lipids, Alabaster, USA) were dissolved 1:1 v/v in CH_3_OH/CHCl_3_ solvent mixture. The solvents were removed by purging N_2_ gas followed by vacuum overnight. The dried lipids were resuspended in 10 mM Tris buffer (pH 7.4) containing 100 mM NaCl, and liposomes were made using the freeze-thaw technique (3-5 cycles). Then, the liposomes were solubilized by mixing them with pentyl-inulin stock solution (100 mg/mL) at 1:1 w/w polymer:lipid ratio and incubating the sample overnight at 4°C under gentle mixing. (*Note*: In the case of incomplete solubilization, 2-3 additional freeze-thaw cycles were performed to achieve complete solubilization of lipids.)

### Reconstitution of APP-C99 into polymer nanodiscs

The solutions of 7:3 w/w DMPC:DMPG polymer nanodiscs, the detergent-purified APP-C99, and the Bio-beads SM2 resin (Bio-Rad) were mixed and incubated at 4 °C for ∼4 h under a slow/careful mixing using a magnetic bar. A 1:30 (w/w) detergent to Bio-beads ratio was used to remove the detergent for the APP-C99 reconstitution. The APP-C99-reconstituted nanodiscs were separated from Bio-beads by filtration. The sample was then purified by size-exclusion chromatography, and the fractions were analyzed by SDS-PAGE. The fractions containing the pure reconstituted protein were used for further characterization.

### Mass spectrometry

The molecular mass of the APP-C99 protein was determined by Matrix-assisted laser desorption/ionization time-of-flight mass spectrometry (MALDI-TOF-MS). One µL of APP-C99 sample was pipetted on 384 well MTP stainless-steel plate followed by adding and mixing one µL of sinapic acid, α□Cyano□4□hydroxycinnamic acid (CHCA) or 2,5-dihydroxybenzoic acid (DHB) matrix. The samples were allowed to air-dry at room temperature before spectra were recorded on Bruker Autoflex mass spectrometer under linear positive ion mode at a voltage of ∼19 kV. The instrument was calibrated in the mid-mass range, 5734.52 Da to 16952.31 Da, using insulin and myoglobin. The spectra were smoothened and corrected for baseline using FlexAnalysis version 3.4 software.

### NMR spectroscopy

All solution NMR experiments were performed on a Bruker 500 MHz NMR spectrometer (Billerica, MA, USA). One-dimensional ^1^H NMR spectra were rec-orded from the SEC-purified APP-C99 reconstituted in nanodiscs. The NMR data were processed and analyzed using Bruker TopSpin (version - 3.6.2) software.

^15^N isotope-labeled APP-C99 was also prepared and reconstituted in polymer nanodiscs. APP-C99 was concentrated by lyophilization and hydrated with water to a final volume of 30 µL (150 µM) and packed into a 3.2 mm NMR rotor. Two-dimensional (2D) [^1^H-^15^N]-HETCOR NMR spectra were recorded using 512 scans and 96 increments (*t*1 dimension) and 2 s recycle delay. NMR data were collected on a 700SB MHz solid-state NMR spectrometer equipped with a 3.2 mm Efree HCN probe and facilitated with a VTN-type variable temperature unit. RF field strengths of 45 kHz on ^1^H and 42.61 kHz on ^15^N, and a 31.82 kHz frequency offset during CP on ^1^H were used. The spectra were recorded at four different temperatures: 243, 253, 273, and 293 K. The success of the experiments is facilitated by the magic-angle spinning (MAS) technique, with a 12.5 kHz MAS frequency. A 0.5 ms cross-polarization (CP) contact time was used to transfer magnetization transfer from proton to nitrogen. Chemical shifts were calibrated using MLF tripeptide as a reference (54).

### Dynamic light scattering (DLS)

DLS measurements were carried out using a Wyatt Technology DynaPro NanoStar instrument equipped with a laser emitting at ∼662 nm. 1 µL of the SEC-purified nanodisc sample was loaded into the quartz MicroCuvette (Wyatt Technology, CA, USA). 10 acquisitions, each 5 s were averaged to obtain the DLS profile. All data were recorded at 25 °C, allowing for at least 10 min for the loaded sample temperature to equilibrate before data acquisition. The spectra were obtained in mass-percentage mode.

### Negative stain transmission electron microscopy (TEM)

The carbon-coated copper 400 mesh TEM grids (Ted Pella Inc., 01702-F) were activated by glow discharge for 45 s using the LEICA EM ACE600 SPUTTER COATER instrument. Five µL of APP nanodiscs sample was adsorbed to the grids for 5 minutes at RT. The grids were washed twice with the Milli-Q and stained with five µL 1% (w/v) uranyl acetate (Ted Pella Inc.). Excess of the uranyl acetate was blotted with the filter paper, and the grids were allowed to air-dry for ∼5 min. Grids with nanodiscs were imaged at a 120 kV accelerating voltage in a JEOL JEM 1400 PLUS electron microscope equipped with a NANOSPRT12 camera at a magnification of 50000X.

### Differential scanning calorimetry

DSC experiments were performed on a differential scanning calorimeter (DSC Nano, TA instruments, New Castle, DE, USA) at a 1 °C/min scan rate under a constant pressure of 3 atm. Heating curves were obtained between the temperature range of 0 °C to 50 °C. The phase transition characterization was investigated for the 7:3 w/w DMPC:DMPG liposomes, protein-free polymer nanodiscs, and polymer nanodiscs reconstituted with APP-C99. The buffer alone was used to obtain the reference DSC profile and was subtracted from each sample type using NanoAnalyze software. The buffers were filtered and degassed before use.

## RESULTS AND DISCUSSION

The APP protein sequences used in this study are shown in Figure 1. C99 is the 99 amino acids containing membrane-anchored region of the full-length APP. It has a C-terminal 6xHis tag flanked by a 16-residues linker sequence (QGRILQISITLAAALE) (**Fig. 1**). The APP-NT2 is the same C99 construct but with an N-terminal 10 His-tag flanked by a linker sequence.

**Figure 1.**
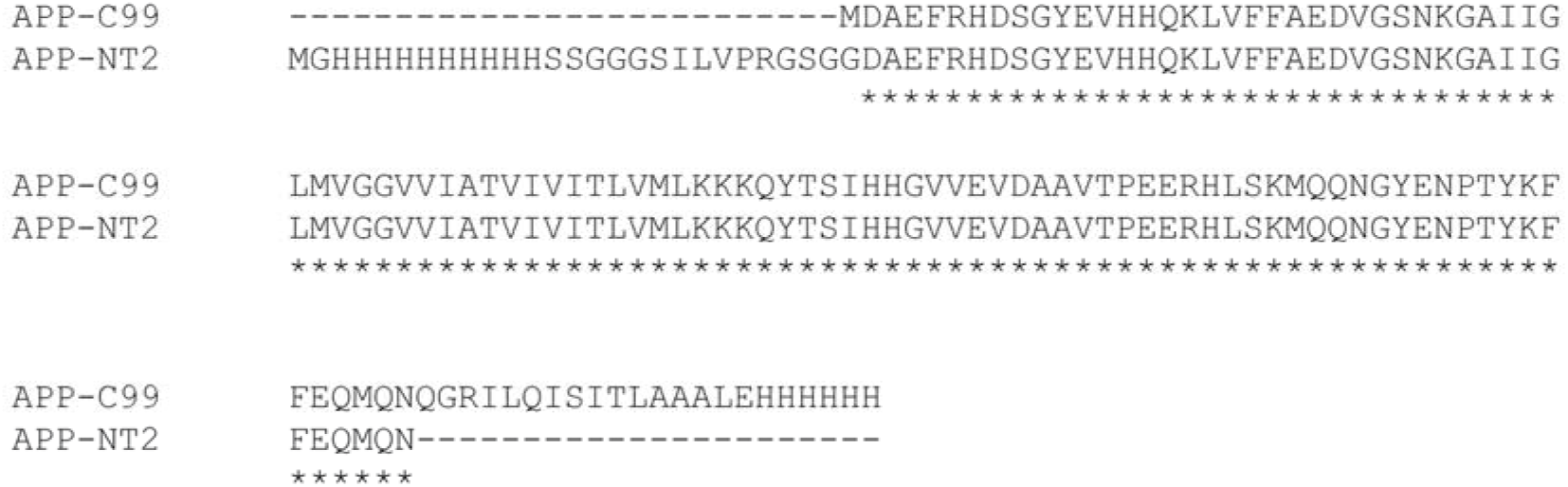
Amino acid sequences of APP-C99 and APP-NT2. “^*^” indicates the conserved residues. APP-NT2 has a 10His-tag followed by a 14-residue-long linker sequence at the N-terminus. APP-C99 has a 6His-tag preceded by a 16-residue long lin quence at the C-terminus.

### Temperature-dependent solubilization of APP-enriched *E. coli* cell membranes

The solubilization was performed at three different temperatures: 4, 25 and 37 °C overnight. The protein band corresponding to APP-NT2 (∼13.9 kDa) was observed near 20-kDa size on the SDS-PAGE gel (**Fig. 2A**). The unusual migration of the APP-NT2 protein on SDS-PAGE gel might be due to the interference caused by polymer and lipids present in the sample. Such unexpected migration of proteins is generally observed for some membrane proteins and also for highly hydrophilic (intrinsically disordered) proteins (55). Another study reported that change in the migration of the protein was due to differential interactions between protein and detergent (SDS), rather than between proteins (56), which recommended other methods together with SDS-PAGE to confirm the size of the protein as well. Overall, the appearance of the APP-NT2 band in the soluble fraction indicates the successful solubilization of APP-NT2-enriched membranes. Due to many *E. coli* protein bands overlapping with the target protein APP-NT2, it was difficult to resolve and visualize the APP-NT2 bands from solubilization at different temperatures. Therefore, the solubilized fractions containing APP-NT2 were further verified by Western-blot analysis.

**Figure 2.**
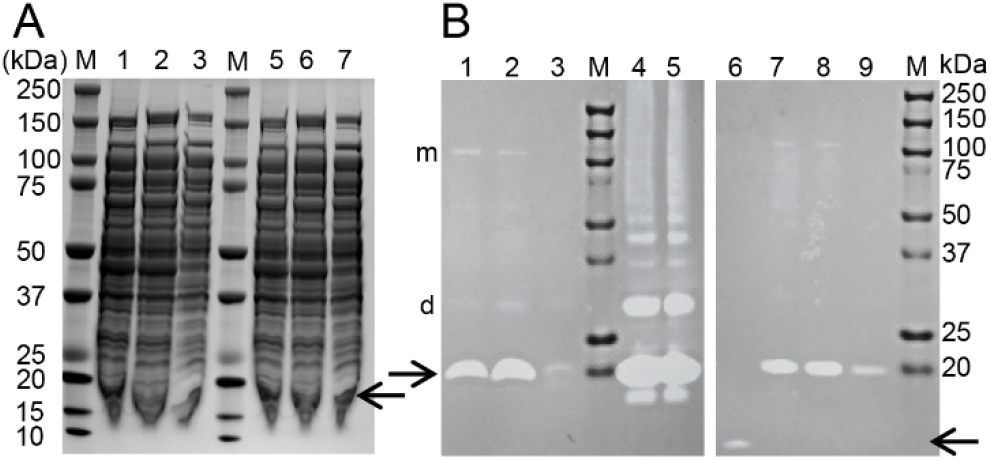
(**A**) Solubilization and SDS-PAGE analysis of APP-NT2-enriched *E. coli* cell membranes. M: protein molecular weight marker, Lanes; (1-3) and (5-7): APP-NT2-enriched membranes solubilized with pentyl-inulin at three different temperatures 4 °C (lanes; 1, 5), 25 °C (lanes; 2, 6) and 37 °C (lanes; 3, 7) (technical replicates). The protein band corresponding to the APP-NT2 is indicated with an arrow. (**B**) Western blot analysis of APP-NT2 in polymer and detergentsolubilized *E. coli* membranes. Lanes 1, 2 and 3 are the mem-branes solubilized by polymer at 4, 25 and 37 °C, respectively. Lanes 7, 8 and 9 are the membranes solubilized by DDM at 4, 25 and 37 °C, respectively. Lanes 4 and 5 were loaded with the insolubilized membrane components from DDM and polymer-solubilized samples respectively. Lane 6: positive control; amyloid β (1-42). M; protein molecular weight marker. The APP-NT2 and Aβ (1-42) bands are indicated with arrows. Protein bands corresponding to dimer and multimer are indicated with d and m, respectively.

### Western blot confirmed the presence of APP-NT2 in polymer-solubilized cell membranes

Western blot was performed on pentyl-inulin solubilized, APP-NT2-enriched *E. coli* cell membranes. 6E10 monoclonal antibody that is specific to the EFRHDS (3-8) sequence motif present within the N-terminal region of Aβ(1-16) was used to detect APP-NT2 in the polymer-solubilized *E. coli* cell membranes. Freshly dissolved Aβ(1-42) was used as a positive control which showed a clear band in the low-molecular-weight region as expected (**Figs. 2B, S1**). The major protein band corresponding to the APP-NT2 was detected in all three samples prepared at three different temperatures (**Figs. 2B, S1**). However, there was a clear difference in the intensity of protein band between the three samples. The sample solubilized at 25 °C showed a strong band followed by 4 °C and then 37 °C, indicating that the pentyl-inulin solubilized APP-NT2-rich *E. coli* membranes more effectively at 25 °C than 4 °C or 37 °C (**Figs. 2B, S1**). Interestingly, the high temperature is not suitable for the solubilization of APP-enriched membranes by either pentylinulin or detergent (DDM). In addition, the intensity of protein bands from polymer samples was higher compared to the DDM-solubilized samples prepared at the same temperature conditions. These observations indicate that pentyl-inulin is more efficient in solubilizing APP-NT2-rich *E. coli* membranes than the commonly used DDM detergent. Strong pro-tein bands were also observed in the insoluble fractions of polymer-solubilized samples. Several low-intensity bands detected by 6E10 were also observed in the high-molecular-weight region indicating some of the APP-NT2 was self-assembled into SDS-resistant dimeric and oligomeric species (**Figs. 2B, S1**). Since APP is known to aggregate and oligomerize, it has been reported to show multiple bands on the gel (57, 58) as reported for Aβ aggregation/oligomerization by in-vitro biophysical/biochemical studies (59) and MD simulations (60). These bands are more prominent in the polymer-solubilized samples compared to the DDM-solubilized samples (**Fig. 2A**). The result also indicates the feasibility of pentyl-inulin for isolating challenging membrane-binding proteins like APP. Further optimization of solubilizing factors such as pH, metal ions, salt and polymer concentration could be helpful to increase the yield of solubilized APP-C99 (53).

### Detergent-solubilization and purification of APP-C99 using Ni-NTA affinity chromatography

The *E. coli* membranes were solubilized overnight using a 20 mM Tris pH 7.8, 150 mM NaCl, 8M urea, and 0.2% SDS. The APP-C99 was purified using Ni-NTA affinity chromatography. The protein binding and the purification were done at room temperature. Prior to elution, the column was washed with 10 column volumes of imidazole-free buffer containing Empigen. Then, the protein was eluted using different concentrations of imidazole. Most of APP-C99 eluted at imidazole concentrations higher than 80 mM (**Fig. S2**). In addition, the impurities were minimal in these fractions. The protein band corresponding to APP-C99 was observed at a slightly higher molecular weight region (>15 kDa) than expected (13.78 kDa) (**Fig. S2**). Such unusual migration of protein on SDS-PAGE gels has been reported in the case of intrinsically disordered proteins (61-63).

### APP-C99 is successfully reconstituted into polymer nanodiscs

7:3 w/w DMPC:DMPG lipid ratio/composition was used to prepare stable non-ionic polymer nanodiscs (64). The APP-C99 fractions from Ni^2+^-NTA purification with minimum impurities were used for nanodisc reconstitution. The detergent (Empigen) was gradually removed using Bio-beads and the reconstituted APP-C99 in polymer nanodiscs was purified by size-exclusion chromatography. A major peak between 37 – 59 mL and a low-intensity peak between 100 – 123 mL were observed (**Fig. 3A**). The fractions from the major peak were analyzed by SDS-PAGE. All the fractions from the major peak showed a single pure protein band (**Fig. 3B**). In the case of the polymer-solubilized sample, multiple protein bands were observed due to oligomerization of APP-C99 (**Fig. 3B**). But a single protein band observed for the detergent-purified and nanodiscs reconstituted APP-C99 sample indicates that inhibition or destabilization of self-assembled APP-C99 due to detergent exposure during purification and stabilization of the monomeric form of APP-C99 in nanodiscs.

**Figure 3.**
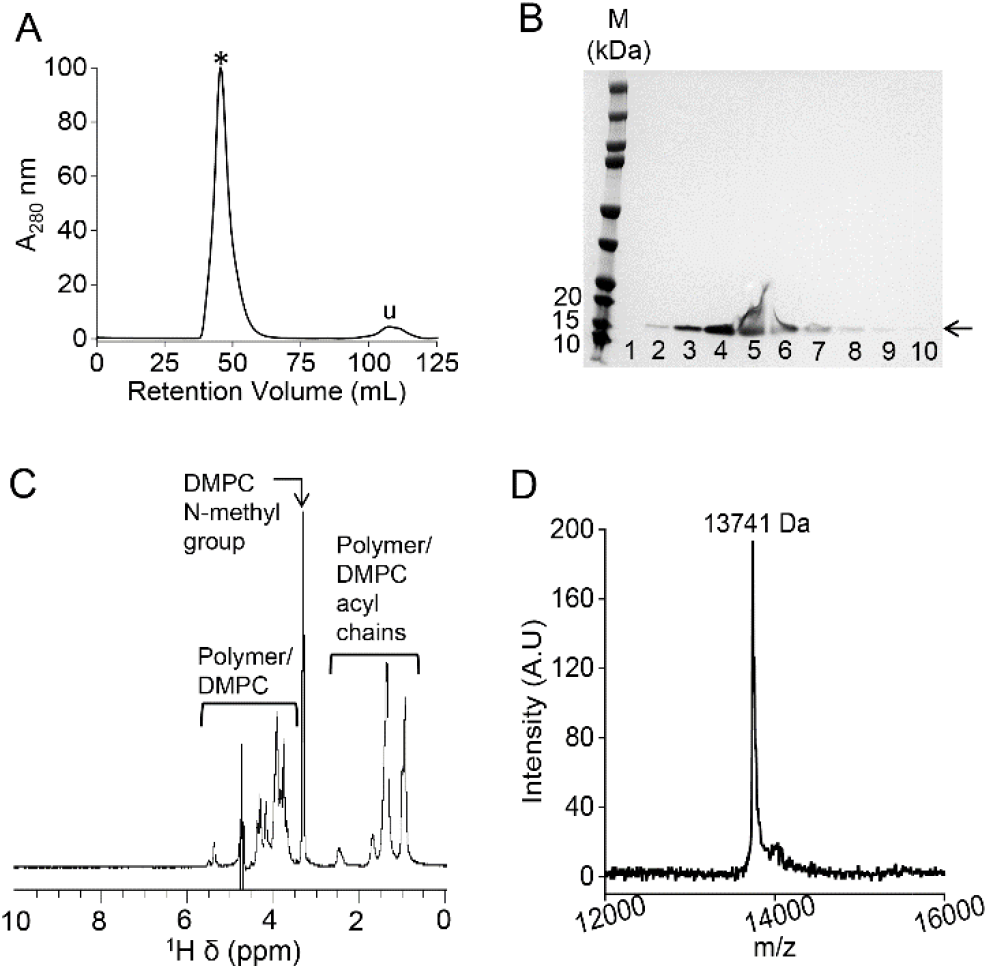
(**A**) Size-exclusion chromatography of nanodisc-reconstituted APP-C99. The elution peak corresponding to APP-C99 is indicated with a ^*^. (**B**) SDS-PAGE analysis of the nanodisc-reconstituted APP-C99 fractions collected from size-exclusion chromatography. The protein band corresponding to APP-C99 is indicated with an arrow. M denotes the protein marker. (**C**) ^1^H NMR spectrum of polymer nanodiscs reconstituted with APP-C99. The protein peaks are not prominent due to broadening caused by lipid-binding in nanodiscs. (**D**) MALDI-TOFF spectrum of APP-C99 reconstituted polymer nanodiscs.

### ^1^H NMR and mass spectrometry

In the ^1^H NMR spectrum, peaks corresponding to the lipids and polymer were observed (**Fig. 3C**) but peaks from APP-C99 were not observed. The absence of protein signals is most likely due to line-broadening caused by the large size of nanodiscs and the interaction of APP-C99 with DMPC/DMPG lipids in polymer nanodiscs. Mass spectrometry showed a major protein peak with 13741 Da which is 30-Da less than the expected mass (13771 Da; (PepCalc)) (**Fig. 3D**). A low-intensity peak at ∼14000 Da was also observed. The mass-to-charge ratio was acquired in a positive ion mode, and the observed high molecular mass species might be due to the formation of alkali metal ion adducts that are reported to occur with the CHCA matrix in a positive ion mode (65). The observed higher mass could be due to strong lipid binding to APP-C99.

### Characterization by DLS, TEM and DSC

The APP-C99 reconstituted polymer nanodiscs were further characterized using DLS and TEM for their size and size/shape respectively. The measured hydrodynamic radius of APP nanodiscs using DLS was ∼9±2 nm, indicating large-size particles present in solution (**Fig. 4A**). This observation was in agreement with the early elution of polymer nanodiscs in size-exclusion chroma-tography. Furthermore, TEM images indicated the nanodisc size as 20±3 nm in diameter (**Fig. 4B**). The morphology of the nanodiscs in the TEM image was nearly spherical, indicating that the integrity of polymer nanodiscs was not affected by APP-C99 reconstitution.

**Figure 4.**
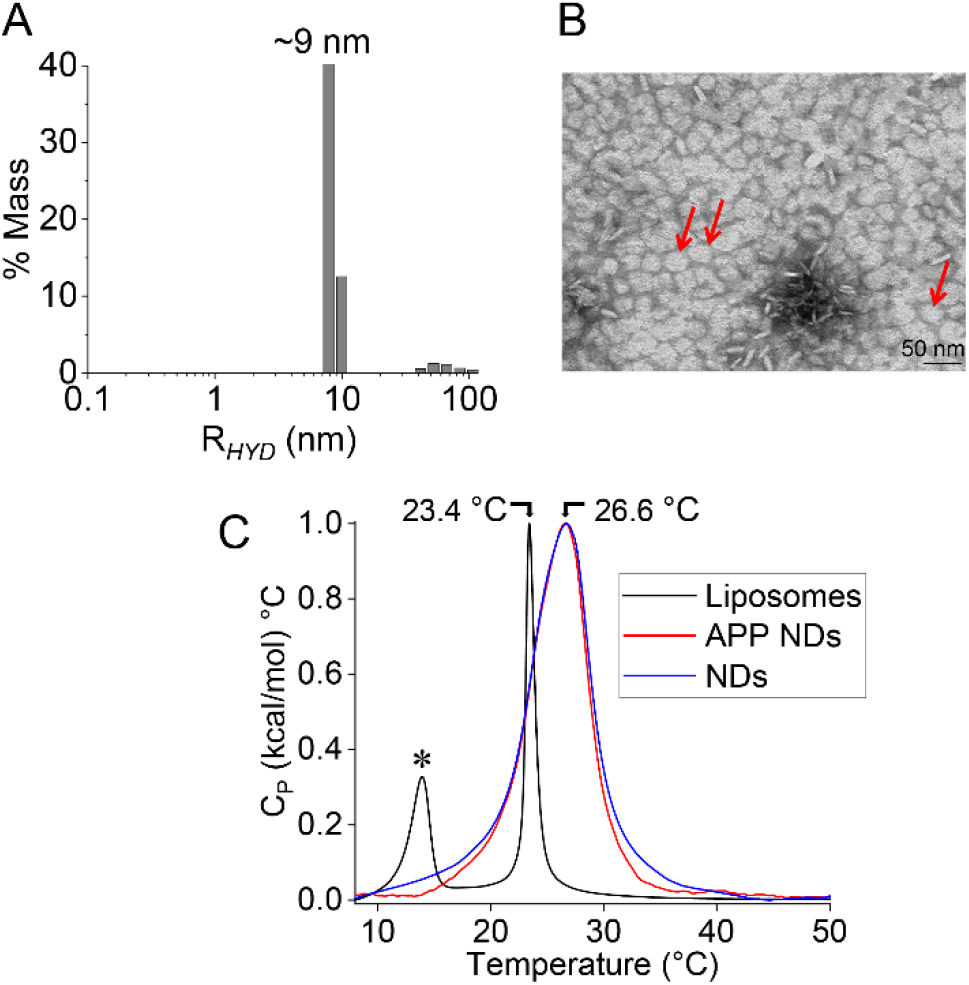
APP-C99 reconstituted 7:3 w/w DMPC:DMPG polymer nanodiscs analyzed by (**A**) DLS, (**B**) TEM, and (**C**) DSC measurements. Reference DSC profiles were recorded using 7:3 w/w DMPC:DMPG liposomes (C; black) and 7:3 w/w DMPC:DMPG polymer nanodiscs (C; blue)and compared with C99-reconstituted 7:3 w/w DMPC:DMPG polymer nanodiscs (C; red). TEM image shows face-on and edge-on (bottom) sides of nanodiscs (∼20 nm diameter). The well-resolved nanodiscs used to measure the nanodisc diameter are indicated with red arrows. The pre-transition peak (13.9 °C) observed for the liposomes in DSC profile is indicated with a ‘LJ’.

The samples were also analyzed by DSC to see whether APP has any effect on the phase transition temperature of lipids in polymer nanodiscs (**Fig. 4C**). The observed T_m_ for the DMPC liposomes (reference) was 23.4 °C. In contrast, the T_m_ for the 7:3 (w/w) DMPC/DMPG in polymer nanodiscs reconstituted with APP-C99 was 26.6 °C, which is 3.2 °C higher than that observed for DMPC liposomes. The T_m_ for the protein-free nanodiscs was similar to that of protein-reconstituted nanodiscs (**Fig. 4C**), indicating the C99 binding did not substantially affect the 7:3 (w/w) DMPC/DMPG lipid order in polymer nanodiscs. The pre-transition/ripple phase peak observed at 14 °C for the DMPC liposomes was not observed for the APP-C99 reconstituted polymer nanodiscs (**Fig. 4C**). The absence of a pre-transition peak was due to the stable bilayer arrangement of lipids in polymer nanodiscs restricting the tilting of DMPC acyl chains. In addition, the phase-transition peak of polymer nanodiscs with and without APP-C99 were broadened substantially than that observed for the 7:3 (w/w) DMPC/DMPG liposomes. The line-broadening and a higher T_m_ observed for the polymer nanodiscs with and without APP-C99 indicate a non-cooperative behavior of lipids due to increased order facilitated by polymer interaction as a stabilizing lipid bilayer belt. Moreover, the interaction of APP with DMPC may also be attributed to the change in the physical phase behavior of DMPC lipids in polymer nanodiscs. However, no substantial difference was observed in the Tm of polymer nanodiscs with and without APP-C99 (**Fig. 4C**).

### Polymer nanodiscs are feasible for high-resolution solid-state NMR studies of APP-C99

Previously, the APP was characterized in lyso-myristoylphosphatidylglycerol (LMPG) detergent micelles using solution NMR spectroscopy (66). MD simulations suggests micelles influence the helical conformation of APP (67). The dimer with the right-handed coiled-coil conformation is dominant in 1-palmitoyl-2-oleoylphosphatidylcholine (POPC) bilayers whereas the dimer with a left-handed conformation was observed in dodecylphosphocholine (DPC) micelles (68, 69). These observations signifying the importance of native membrane environment for the structural studies of membrane proteins in general. In this study, we wanted to examine how the reconstitution into lipid-bilayer affects the quality of NMR spectrum. 150 µM ^15^N-isotope labeled APP-C99 was reconstituted in 7:3 (w/w) DMPC/DMPG polymer nanodiscs and characterized by solid-state NMR spectroscopy under MAS conditions (**Fig. 5**). Since the amount of protein (150 µM) present in the **30** µL nanodisc sample in 3.2 mm rotor is very small, it is a challenge to obtain high-resolution NMR spectra for APP-C99. The 2D spectrum displays well resolved ^15^N-^1^H resonances. By using a short CP contact time, resonances from immobile residues of the protein that possess strong ^1^H-^15^N dipolar couplings can be selectively observed. On the other hand, residues from the soluble domain of the protein are expected to undergo faster motion as compared to transmembrane/membrane-bound residues and therefore require a longer CP contact time as they are associated with weak motionally-averaged ^1^**H**-^15^N dipolar couplings. Therefore, with 0.5 s CP contact time used in acquiring the 2D spectrum, most of the observed resonances are expected to be from the lipid bilayer-embedded amino acid residues and those making strong contacts with the lipid bilayer surface of nanodisc. However, due to extensive overlap of resonances, only few NMR assignments were made based on the chemical shifts reported in literature (66). The spectral broadening observed in the current study is substantially large compared to that reported in micelles/bicelles (66). Peak overlap became severe at lower temperature (243 K -273 K) likely due to the less dynamic gel phase of lipids (**Fig. S3**). The increased peak broadening observed in nanodiscs is due to large size (∼20 nm diameter) of nanodiscs and also due to the low temperature conditions (273 – 243 K) used to record NMR spectra. Overall, the NMR data suggests the feasibility of using high-resolution solid-state NMR to characterize APP-C99 in polymer nanodiscs in future studies. Further enhancement in resolution can be achieved by using deuterated lipids and faster MAS experiments. Assignment of resonances and structure determination can be accomplished by employing 3D MAS experiments and using ^13^C-^15^N-labeled protein and deuterated lipids as needed.

**Figure 5.**
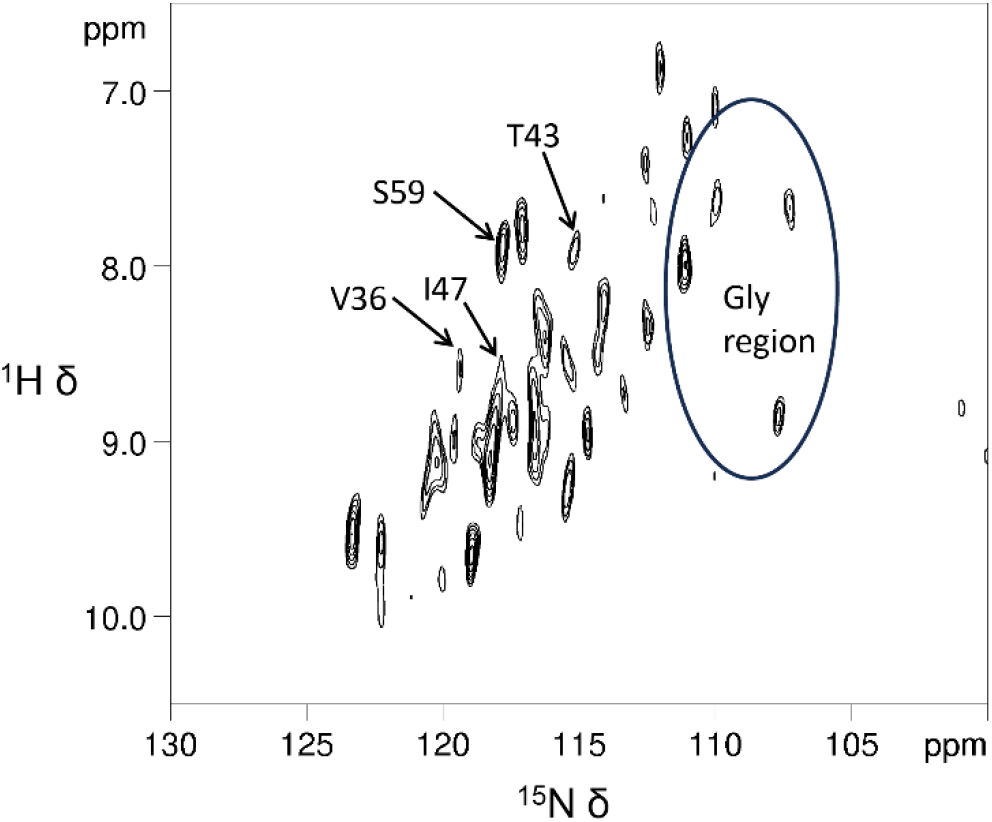
2D [^1^H-^15^N]-HETCOR NMR spectrum of APP-C99 reconstituted into 7:3 w/w DMPC:DMPG polymer nanodiscs. The spectrum was recorded on a 700 MHz Bruker NMR spectrometer at 293 K. A 0.5 s CP contact time was used to detect lipid embedded/bound amino acid residues of APP-C99. Partial assignments are shown. The 30 µL of 150 µM APP-C99 reconstituted in 7:3 (w/w) DMPC/DMPG polymer nanodiscs, 10 mM potassium phosphate buffer (pH 7.4) was packed into 3.2 mm MAS NMR rotor.

## CONCLUSIONS

In this study, we have successfully demonstrated the direct isolation and reconstitution of APP-C99 in pentyl-inulin polymer based nanodiscs. We have also successfully reconstituted the detergent-purified APP-C99 in 7:3 (w/w) DMPC/DMPG pentyl-inulin polymer nanodiscs. The APP-C99 reconstituted nanodiscs were characterized by a combination of SEC, DLS and TEM measurements that revealed stable and homogeneous size nanodiscs. The presence of APP-C99 did not substantially influence the physical behaviour of lipids encased in the polymer nanodiscs. Moreover, the feasibility of isolating APP-C99 in native lipids using polymer is demonstrated. Room temperature is shown to be more effective for the polymer-based solubilization of APP-C99-enriched bacterial cell membranes. Solid-state NMR spectra acquired under MAS conditions indicated the feasibility of using polymer nanodiscs for high-resolution structural and dynamical studies of APP-C99.

## ASSOCIATED CONTENT

### Supporting Information

The Supporting Information is available free of charge on the ACS Publications website.

Western blot, SDS-PAGE and 2D NMR.

## AUTHOR INFORMATION

### Authors

**Bankala Krishnarjuna - Biophysics Program, Department of Chemistry, Macromolecular Science and Engineering, Biomedical Engineering, University of Michigan, Ann Arbor, MI 48109, USA**.

**Gaurav Sharma - Biophysics Program, Department of Chemistry, Macromolecular Science and Engineering, Biomedical Engineering, University of Michigan, Ann Arbor, MI 48109, USA**.

**Volodymyr M Hiiuk - Biophysics Program, Department of Chemistry, Macromolecular Science and Engineering, Biomedical Engineering, University of Michigan, Ann Arbor, MI 48109, USA**.

**Jochem Struppe - Bruker Biospin Corporation, 15 Fortune Drive, Billerica, Massachusetts 01821, USA**.

**Pavel Nagorny - Department of Chemistry, University of Michigan, Ann Arbor, Michigan 48109, USA**.

**Magdalena I Ivanova - Department of Neurology, University of Michigan, Ann Arbor, Michigan 48109, USA**

### Notes

The authors declare no competing financial interest.

## ACKNOWLEDGMENT

This study was supported by National Institutes of Health (NIH) (R35 GM139572 and RO1 DK132214 to A.R. and R35 GM136341 to P. N.). We thank Dr. Thirupathi Ravula for help with the synthesis of pentyl-inulin polymer used in this study and Professor Charles Sanders from the Department of Biochemistry, Vanderbilt University School of Medicine for kindly providing APP plasmid constructs.

## REFERENCES

(1) Gulezian, E.; et al. Membrane protein production and formulation for drug discovery. Trends Pharmacol. Sci. 2021, 42, 657–674.

(2) Levental, I.; Lyman, E. Regulation of membrane protein structure and function by their lipid nano-environment. Nat. Rev. Mol. Cell Biol. 2023, 24, 107–122.

(3) Piper, S. J.; et al. Membranes under the magnetic lens: A dive into the diverse world of membrane protein structures using Cryo-EM. Chem. Rev. 2022, 122, 13989–14017.

(4) Overington, J. P.; et al. How many drug targets are there? Nat. Rev. Drug Discov. 2006, 5, 993–996.

(5) Krishnarjuna, B.; Ramamoorthy, A. Detergent-free isolation of membrane proteins and strategies to study them in a near-native membrane environment. Biomolecules 2022, 12, 1076.

(6) Young, J. W. Recent advances in membrane mimetics for membrane protein research. Biochem. Soc. Trans. 2023, 51, 1405–1416.

(7) Bayburt, T. H.; et al. SelfUassembly of discoidal phospholipid bilayer nanoparticles with membrane scaffold proteins. Nano Lett. 2002, 2, 853–856.

(8) Ritchie, T. K.; et al. Reconstitution of membrane proteins in phospholipid bilayer nanodiscs. Methods Enzymol. 2009, 464, 211–231.

(9) Bayburt, T. H.; Sligar, S. G. Membrane protein assembly into nanodiscs. FEBS Lett. 2010, 584, 1721–1727.

(10) Shen, H. H.; et al. Reconstitution of membrane proteins into model membranes: seeking better ways to retain protein activities. Int. J. Mol. Sci. 2013, 14, 1589–1607.

(11) Yeung, Y.-G.; Stanley, E. R. Rapid detergent removal from peptide samples with ethyl acetate for mass spectrometry analysis. Curr. Protoc. Protein Sci. 2010, 59, 16.12.11-16.12.15.

(12) Bao, H.; et al. The maltose ABC transporter: Action of membrane lipids on the transporter stability, coupling and ATPase activity. BBA-Biomembranes 2013, 1828, 1723–1730.

(13) Yang, Z.; et al. Membrane protein stability can be compromised by detergent interactions with the extramembranous soluble domains. Protein Sci. 2014, 23, 769–789.

(14) Seddon, A. M.; et al. Membrane proteins, lipids and detergents: not just a soap opera. BBA-Biomembranes 2004, 1666, 105–117.

(15) Lambert, N. A. GPCR dimers fall apart. Science Signaling 2010, 3, pe12–pe12.

(16) Marty, M. T.; et al. Interfacing membrane mimetics with mass spectrometry. Accounts of Chemical Research 2016, 49, 2459–2467.

(17) Young, J. W.; et al. Development of a method combining peptidiscs and proteomics to identify, stabilize, and purify a detergent-sensitive membrane protein assembly. Journal of Proteome Research 2022, 21, 1748–1758.

(18) Dürr, U. H. N.; et al. The magic of bicelles lights up membrane protein structure. Chem. Rev. 2012, 112, 6054–6074.

(19) Lucyanna, B.-B.; et al. Structural versatility of bicellar systems and their possibilities as colloidal carriers. In Pharmaceutics, 2011; Vol. 3, pp 636–664.

(20) Majeed, S.; et al. Lipid membrane mimetics in functional and structural studies of integral membrane proteins. In Membranes, 2021; Vol. 11.

(21) Thoma, J.; Burmann, B. M. Fake it ‘till you make it—the pursuit of suitable membrane mimetics for membrane protein biophysics. In Int. J. Mol. Sci., 2021; Vol. 22.

(22) Christian Isalomboto, N.; et al. General perception of liposomes: formation, manufacturing and applications. Liposomes-Advances and Perspectives 2019, Ch. 3.

(23) Zhang, J. X.; et al. Application of liposomes in drug development--focus on gastroenterological targets. Int. J. Nanomed. 2013, 8, 1325–1334.

(24) Carlson, M. L.; et al. The Peptidisc, a simple method for stabilizing membrane proteins in detergent-free solution. eLife 2018, 7, e34085.

(25) Cross, T. A.; et al. Influence of solubilizing environments on membrane protein structures. Trends Biochem. Sci. 2011, 36, 117–125.

(26) Das, B. B.; et al. Structure determination of a membrane protein in proteoliposomes. J. Am. Chem. Soc. 2012, 134, 2047–2056.

(27) Kijac, A. Z.; et al. Magic-angle spinning solid-state NMR spectroscopy of nanodisc-embedded human CYP3A4. Biochemistry 2007, 46, 13696–13703.

(28) Luthra, A.; et al. Nanodiscs in the studies of membrane-bound cytochrome P450 enzymes. Methods Mol. Biol. 2013, 987, 115–127.

(29) Hagn, F.; et al. Optimized phospholipid bilayer nanodiscs facilitate high-resolution structure determination of membrane proteins. J. Am. Chem. Soc. 2013, 135, 1919–1925.

(30) Denisov, I. G.; Sligar, S. G. Nanodiscs for structural and functional studies of membrane proteins. Nat. Struct. Mol. Biol. 2016, 23, 481–486.

(31) Denisov, I. G.; Sligar, S. G. Nanodiscs in membrane biochemistry and biophysics. Chem. Rev. 2017, 117, 4669–4713.

(32) Nasr, M. L.; et al. Covalently circularized nanodiscs for studying membrane proteins and viral entry. Nat. Methods 2017, 14, 49–52.

(33) Hagn, F.; et al. Assembly of phospholipid nanodiscs of controlled size for structural studies of membrane proteins by NMR. Nat. Protoc. 2018, 13, 79–98.

(34) Nasr, M. L.; Wagner, G. Covalently circularized nanodiscs; challenges and applications. Curr. Opin. Struct. 2018, 51, 129–134.

(35) Sligar, S. G.; Denisov, I. G. Nanodiscs: A toolkit for membrane protein science. Protein Sci. 2021, 30, 297–315.

(36) Zhang, M.; et al. Cryo-EM structure of an activated GPCR– G protein complex in lipid nanodiscs. Nat. Struct. Mol. Biol. 2021, 28, 258–267.

(37) Janata, M.; et al. Sulfonated polystyrenes: pH and Mg2+ insensitive amphiphilic copolymers for detergent-free membrane protein isolation. Eur. Polym. J. 2023, 198, 112412.

(38) Chen, G. F.; et al. Amyloid beta: structure, biology and structure-based therapeutic development. Acta Pharmacol. Sin. 2017, 38, 1205–1235.

(39) Zheng, H.; Koo, E. H. Biology and pathophysiology of the amyloid precursor protein. Molecular Neurodegeneration 2011, 6, 27.

(40) van der Kant, R.; Goldstein, L. S. Cellular functions of the amyloid precursor protein from development to dementia. Dev. Cell 2015, 32, 502–515.

(41) en, J.; Kelleher, R. J., 3rd The presenilin hypothesis of Alzheimer’s disease: evidence for a loss-of-function pathogenic mechanism. Proc. Natl. Acad. Sci. USA 2007, 104, 403–409.

(42) Beel, A. J.; et al. Direct binding of cholesterol to the amyloid precursor protein: An important interaction in lipid-Alzheimer’s disease relationships? Biochim. Biophys. Acta Mol. Cell Biol. Lipids 2010, 1801, 975–982.

(43) Lane, C. A.; et al. Alzheimer’s disease. European Journal of Neurology 2018, 25, 59–70.

(44) Sehar, U.; et al. Amyloid beta in aging and alzheimer’s disease. Int. J. Mol. Sci. 2022, 23.

(45) Haass, C.; et al. Trafficking and proteolytic processing of APP. CSH Perspect. Med. 2012, 2, a006270.

(46) Capone, R.; et al. The C99 domain of the amyloid precursor protein resides in the disordered membrane phase. J. Biol. Chem. 2021, 296, 100652.

(47) Pfundstein, G.; et al. Amyloid precursor protein (APP) and amyloid β (Aβ) interact with cell adhesion molecules: implications in alzheimer’s disease and normal physiology. Front. Dell Dev. Biol. 2022, 10, 969547.

(48) Ravula, T.; Ramamoorthy, A. Synthesis, characterization, and nanodisc formation of non-ionic polymers. Angew. Chem. Int. Ed. 2021, 60, 16885–16888.

(49) Krishnarjuna, B.; et al. Detergent-free isolation of CYP450-reductase’s FMN-binding domain in E.coli lipid-nanodiscs using a charge-free polymer. ChemComm 2022, 58, 4913–4916.

(50) Krishnarjuna, B.; et al. Non-ionic inulin-based polymer nanodiscs enable functional reconstitution of a redox complex composed of oppositely charged CYP450 and CPR in a lipid bilayer membrane. Anal. Chem. 2022, 94, 11908–11915.

(51) Studier, F. W. Protein production by auto-induction in high density shaking cultures. Protein Expr. Purif. 2005, 41, 207–234.

(52) Bentley, M. R.; et al. Rapid elaboration of fragments into leads by X-ray crystallographic screening of parallel chemical libraries (REFiLX). J. Med. Chem. 2020, 63, 6863–6875.

(53) Krishnarjuna, B.; et al. Factors influencing the detergent-free membrane protein isolation using synthetic nanodisc-forming polymers. BBA-Biomembranes 2024, 1866, 184240.

(54) Kumari, B.; et al. Efficient referencing of FSLG CPMAS HETCOR spectra using 2D 1H–1H MAS FSLG. Appl. Magn. Reson. 2019, 50, 1399–1407.

(55) Rath, A.; et al. Detergent binding explains anomalous SDS-PAGE migration of membrane proteins. Proc. Natl. Acad. Sci. USA 2009, 106, 1760–1765.

(56) Walkenhorst, W. F.; et al. Polar residues in transmembrane helices can decrease electrophoretic mobility in polyacrylamide gels without causing helix dimerization. BBA-Biomembranes 2009, 1788, 1321–1331.

(57) Gerber, H.; et al. Zinc and copper differentially modulate amyloid precursor protein processing by γ-secretase and amyloid-β peptide production. J. Biol. Chem. 2017, 292, 3751–3767.

(58) Perrin, F.; et al. Dimeric transmembrane orientations of APP/C99 regulate γ-secretase processing line impacting signaling and oligomerization. iScience 2020, 23, 101887.

(59) Fawzi, N. L.; et al. Atomic-resolution dynamics on the surface of amyloid-β protofibrils probed by solution NMR. Nature 2011, 480, 268–272.

(60) Okumura, H.; Itoh, S. G. Molecular dynamics simulation studies on the aggregation of amyloid-β peptides and their disaggregation by ultrasonic wave and infrared laser irradiation. Molecules 2022, 27.

(61) MacRaild, C. A.; et al. Conformational dynamics and antigenicity in the disordered malaria antigen merozoite surface protein 2. PLoS One 2015, 10, e0119899.

(62) Morales, R. A. V.; et al. Structural basis for epitope masking and strain specificity of a conserved epitope in an intrinsically disordered malaria vaccine candidate. Sci. Rep. 2015, 5, 10103.

(63) Krishnarjuna, B.; et al. Strain-transcending immune response generated by chimeras of the malaria vaccine candidate merozoite surface protein 2. Sci. Rep. 2016, 6, 20613.

(64) Krishnarjuna, B.; et al. Enhancing the stability and homogeneity of non-ionic polymer nanodiscs by tuning electrostatic interactions. J. Colloid Interface Sci. 2023, 634, 887–896.

(65) Lou, X.; et al. Dual roles of [CHCA + Na/K/Cs]+ as a cation adduct or a protonated salt for analyte ionization in matrix-assisted laser desorption/ionization mass spectrometry. Rapid Commun. Mass Spectrom. 2021, 35, e9111.

(66) Barrett, P. J.; et al. The amyloid precursor protein has a flexible transmembrane domain and binds cholesterol. Science 2012, 336, 1168–1171.

(67) Dominguez, L.; et al. Transmembrane fragment structures of amyloid precursor protein depend on membrane surface curvature. J. Am. Chem. Soc. 2014, 136, 854–857.

(68) Nadezhdin, K. D.; et al. Dimeric structure of transmembrane domain of amyloid precursor protein in micellar environment. FEBS Lett. 2012, 586, 1687–1692.

(69) Dominguez, L.; et al. Impact of membrane lipid composition on the structure and stability of the transmembrane domain of amyloid precursor protein. Proc. Natl. Acad. Sci. USA 2016, 113, E5281-E5287.

