## Supporting Information for "Nanodisc reconstitution and characterization of amyloid-β precursor protein C99"

Ayyalusamy Ramamoorthy

**=**equal contribution

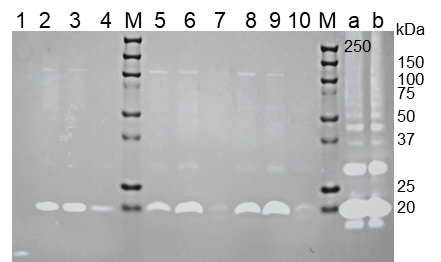

**Figure S1.** Western blot analysis of APP-NT2 in polymer and detergent-solubilized *E. coli* membranes. Lane 1: positive control; amyloid β (1-42). Lanes 2, 3, and 4 are the membranes solubilized by DDM at 4, 25, and 37 °C, respectively. Lanes 5, 6, and 7 (and 8, 9, and 10 (technical replicates)) are the membranes solubilized by polymer at 4, 25, and 37 °C, respectively. Lanes a and b were loaded with the insolubilized membrane components from DDM and polymer-solubilized samples, respectively. M; protein molecular weight marker.

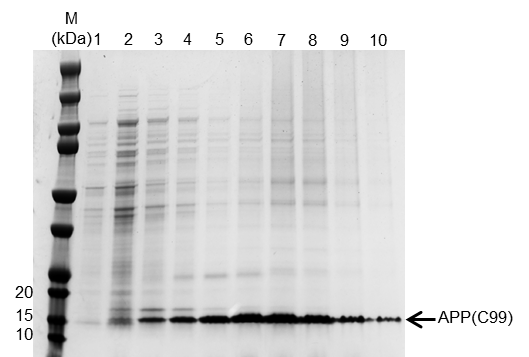

**Figure S2.** Ni^2+^-NTA affinity purification of APP. The 1-10 indicates different concentrations of imidazole used to elute the protein. The protein band corresponding to APP(C99) is labelled. M denotes the protein marker.

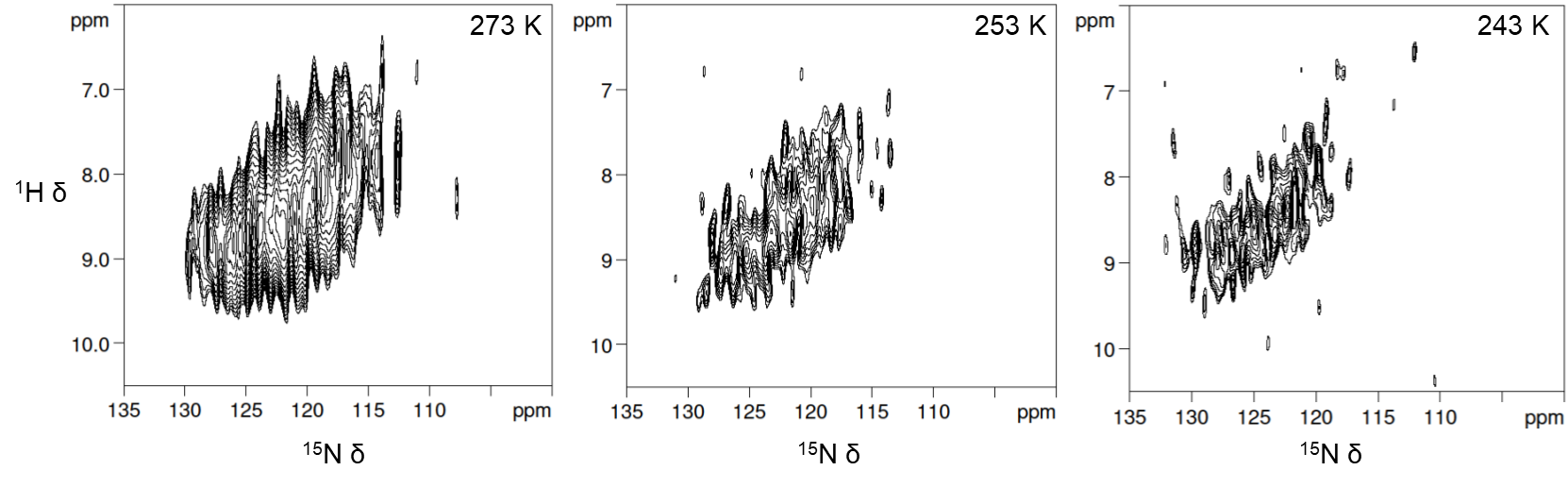

**Figure S3.** 2D [^1^H-^15^N]-HETCOR NMR spectra of APP-C99 reconstituted into 7:3 w/w DMPC:DMPG polymer nanodiscs. The spectra were recorded on a 700 MHz Bruker NMR spectrometer operated with the probe temperature as indicated. 0.5 s CP contact time was used to detect lipid-embedded/bound amino acid residues of APP-C99.
